# Kynurenic acid inflammatory signaling expands in primates and impairs prefrontal cortical cognition

**DOI:** 10.1101/2024.06.13.598842

**Authors:** Shengtao Yang, Dibyadeep Datta, Fenna M. Krienen, Emi Ling, Elizabeth Woo, Athena May, George M. Anderson, Veronica C. Galvin, Guillermo Gonzalez-Burgos, David A. Lewis, Steven A. McCarroll, Amy FT Arnsten, Min Wang

## Abstract

Cognitive deficits from dorsolateral prefrontal cortex (dlPFC) dysfunction are common in neuroinflammatory disorders, including long-COVID, schizophrenia and Alzheimer’s disease, and have been correlated with kynurenine inflammatory signaling. Kynurenine is further metabolized to kynurenic acid (KYNA) in brain, where it blocks NMDA and α7-nicotinic receptors (nic-α7Rs). These receptors are essential for neurotransmission in dlPFC, suggesting that KYNA may cause higher cognitive deficits in these disorders. The current study found that KYNA and its synthetic enzyme, KAT II, have greatly expanded expression in primate dlPFC in both glia and neurons. Local application of KYNA onto dlPFC neurons markedly reduced the delay-related firing needed for working memory via actions at NMDA and nic-α7Rs, while inhibition of KAT II enhanced neuronal firing in aged macaques. Systemic administration of agents that reduce KYNA production similarly improved cognitive performance in aged monkeys, suggesting a therapeutic avenue for the treatment of cognitive deficits in neuroinflammatory disorders.

## INTRODUCTION

Kynurenine signaling is increased under conditions of inflammation, when tryptophan is metabolized by indoleamine 2,3-dioxygenase (IDO) to generate kynurenine instead of serotonin. Although extensive research has examined kynurenine’s roles in the immune response^1^, and apoptotic cell death^2,3^, recent research suggests that it also plays a major role in the cognitive deficits caused by many inflammatory disorders. For example, Post-Acute Sequelae of COVID-19 (PASC, “long-COVID”) is associated with increased kynurenine signaling^4–7^, and the cognitive deficits caused by long-COVID correlate most strongly with plasma kynurenine levels^5^. A similar relationship is seen in patients with schizophrenia^8^, and in aging and Alzheimer’s disease (AD;^9^), where plasma kynurenine levels correlate with cognitive deficits. These data suggest that kynurenine signaling may have significant effects on the higher cortical circuits that subserve cognitive functioning.

The pattern of cognitive deficits in both long-COVID and schizophrenia fits with preferential dysfunction of the dorsolateral prefrontal cortex (dlPFC), the recently evolved cortical region that subserves working memory and higher cognition in primates^10^. For example, many patients with long-COVID report “brain fog” associated with impaired working memory and executive function^11–15^, including impaired recall but not recognition memory, coherent with dysfunction of the dlPFC but not the medial temporal lobe^16^. Multiple brain imaging studies of schizophrenia have established dlPFC underactivity during working memory, e.g. that correlates with thought disorder^17–20^. However, nothing is known about how kynurenine signaling is expressed in dlPFC, or how it impacts dlPFC neuronal function.

Layer III of the dlPFC contains the recurrent excitatory microcircuits essential to working memory and executive function, allowing neurons to excite each other to maintain firing when memory is needed to keep information “in mind”^21^. The dlPFC contains “Delay cells” that are able to sustain firing across the delay period in a working memory task, e.g. with selective response to a location in space during a visuospatial task^22^. Previous research has shown that the ability to maintain firing across the delay depends on both glutamate stimulation of NMDA glutamate receptors (NMDARs) and cholinergic stimulation of α7-nicotinic receptors (nic-α7Rs), with surprisingly little contribution from AMPA glutamate receptors that usually contribute to glutamate actions^23,24^. Both NMDAR and nic-α7R are found within the post-synaptic density (PSD) on layer III dlPFC spines^23,24^, the likely substrate of the recurrent excitatory connections that support delay cell firing. The current study examined whether these circuits are affected by kynurenine signaling.

Increased kynurenine metabolism from tryptophan under inflammatory conditions occurs in both brain and the periphery^25^, where kynurenine is actively taken up into brain from blood, and is also synthesized locally in brain, where rodent studies show expression in glia^26–28^. Kynurenine can be further metabolized to either kynurenic acid (KYNA, also known as kynurenate) by KAT II, encoded by the gene *AADAT* (see graphic abstract), or to quinolinic acid (QUIN) in a parallel pathway^29^. These metabolites are charged, and thus normally do not cross the blood brain barrier, but can be synthesized directly in brain^26^. Longitudinal measures in humans show that kynurenine levels increase, and serotonin levels decrease with advancing age and are associated with increased fraility^30^. There are also age-related increases in KYNA levels in humans and animals^31,32^, including increases in the nonhuman primate cortex^33^. KYNA levels are also increased in the brains of patients who died from SARS-COV-2 infection^34^, and in the dlPFC of patients with schizophrenia, especially those who exhibit an elevated inflammatory profile^8,35^ and/or treatment resistance^36^. IDO levels are also increased near plaques and tangles in AD cortex ^37^, and there is increased KYNA in AD brain^38^. Thus, it is clinically important to learn how KYNA impacts dlPFC physiology and function.

KYNA and QUIN have opposite effects on NMDAR neurotransmission: QUIN stimulates, while KYNA blocks, NMDARs^39^. KYNA has high affinity for the glycine site on the NMDAR^39,40^, with much lower affinity for AMPAR and kainate receptors^40^. Especially relevant to dlPFC neurotransmission, KYNA also blocks nic-α7Rs^41–43^. Based on its NMDAR blocking properties, KYNA has come to be known as the “protective” metabolite, as under extreme conditions of high glutamate release, such as ischemic stroke, or with *in vitro* models, KYNA blockade of NMDAR can protect neurons from rapid death by apoptosis^39^. However, we have hypothesized that KYNA would be harmful under conditions of normal or reduced glutamate release, and would particularly impair the functioning of circuits such as those in the dlPFC that depend on both NMDAR and nic-α7R neurotransmission. This hypothesis would be consistent with dlPFC cognitive deficits being central to many inflammation-related disorders, including long-COVID, schizophrenia, aging and AD, as described above. The current study examined the expression patterns of *AADAT* in mouse, macaque and human PFC, and performed in depth studies of KYNA and KAT II protein expression in aged macaque dlPFC, including ultrastructural localization of KYNA to visualize its relationship to glutamate synapses on spines. We additionally examined the effects of KYNA vs. KAT-II inhibition on dlPFC Delay cell firing, and the effects of systemic administration of KAT-II and IDO inhibitors on working memory function in aged macaques with naturally-occurring reductions in dlPFC neuronal firing^44^ and cognitive deficits^45–49^ beginning in middle age. The results show a large expansion of KYNA signaling in primate dlPFC which markedly reduces the neuronal firing needed for higher cognition, with KAT II inhibition helping to restore cognitive function, suggesting a new avenue for the treatment of cognitive deficits in long-COVID and other neuroinflammatory mental disorders.

## RESULTS

### Transcriptomic expression of AADAT is greater in primates than in mouse

We analyzed single cell RNA sequencing data from mouse frontal cortex^50^, and single nucleus data from the dlPFC of macaque^51^ and human^52^ to investigate the expression of *AADAT,* encoding the KAT II enzyme that synthesizes KYNA, showed marked species differences between mouse mPFC and primate dlPFC. In mouse, there was very low-to-undetectable expression of *AADAT* across all cell types, and it was further concentrated in glia, with limited expression in neurons (Fig. 1a). In contrast, there was much more (∼200x) expression of *AADAT* in macaque dlPFC, where in addition to relatively high expression in astrocytes and oligodendrocytes, there was detectable expression in all neuron types (Fig. 1b). Indeed, the cells with the highest levels of *AADAT* expression were pyramidal cells (layers 3-5) that co-expressed *RORB*, which is found throughout multiple layers in macaque dlPFC, including CUX2-expressing superficial pyramidal cells e.g. in layer III (Fig. S1). There was lower expression in excitatory neuron types that tend to be found in deeper layers (Fig. 1b). Expression of *AADAT* was similar in human dlPFC, with relatively high levels in most pyramidal cell and interneuron subgroups, and in glia (Fig. 1c). Thus, there is an expansion of *AADAT* expression across PFC evolution, especially evident in superficial pyramidal cells (Fig. 1d).

**Figure 1.**
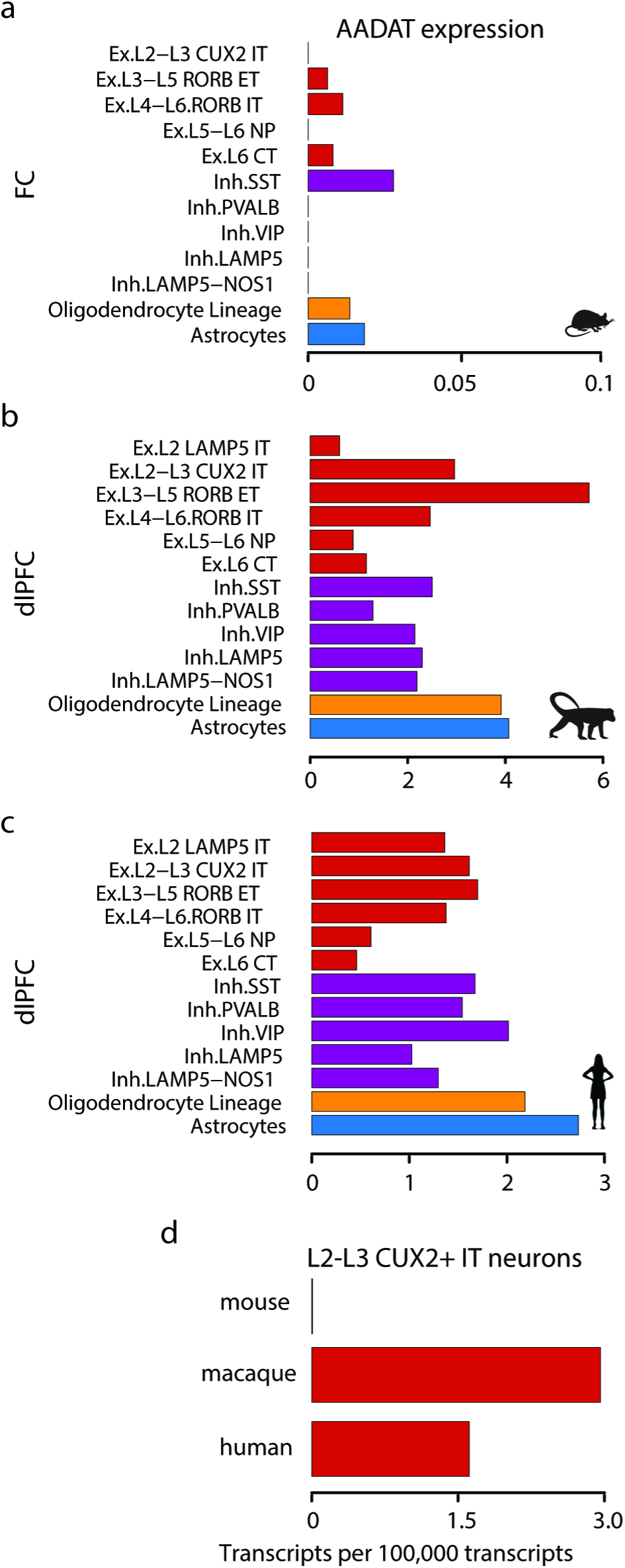
*AADAT* expression in mouse, macaque, and human. *AADAT* encodes KAT II, the enzyme that metabolizes kynurenine into kynurenic acid. Single cell/nucleus RNA expression data are from frontal cortex in mouse^50^ **(a)**, and from dlPFC in macaque^51^ **(b)** and human^52^ **(c)**. Note the large differences in scale between mouse and primates, where normalized levels in mouse are much lower, and more focused in glia than in primates. The differences between mouse and primates were noteworthy for the layer 2-3 CUX2-expressing excitatory neurons **(d)**.

### KAT II and KYNA are in neurons as well as astrocytes in macaque dlPFC

Cellular localization of KAT II-Protein labeling of KAT II in the macaque dlPFC was consistent with the transcriptomic profiles, showing KAT II expression in neurons as well as astrocytes in aged macaque layer III dlPFC. Multiple label immunofluorescence (MLIF) showed that KAT II was expressed in pyramidal cells and astrocytes, co-labeled by MAP2 (Figs. 2a,b) and GFAP (Figs. 2c,d), respectively. Some of the labeled astrocytes showed a reactive-like phenotype (e.g. Fig. 2c) and were in contact with blood vessels (Fig. 2d) or neuronal cell bodies (Fig. 2c).

**Figure 2.**
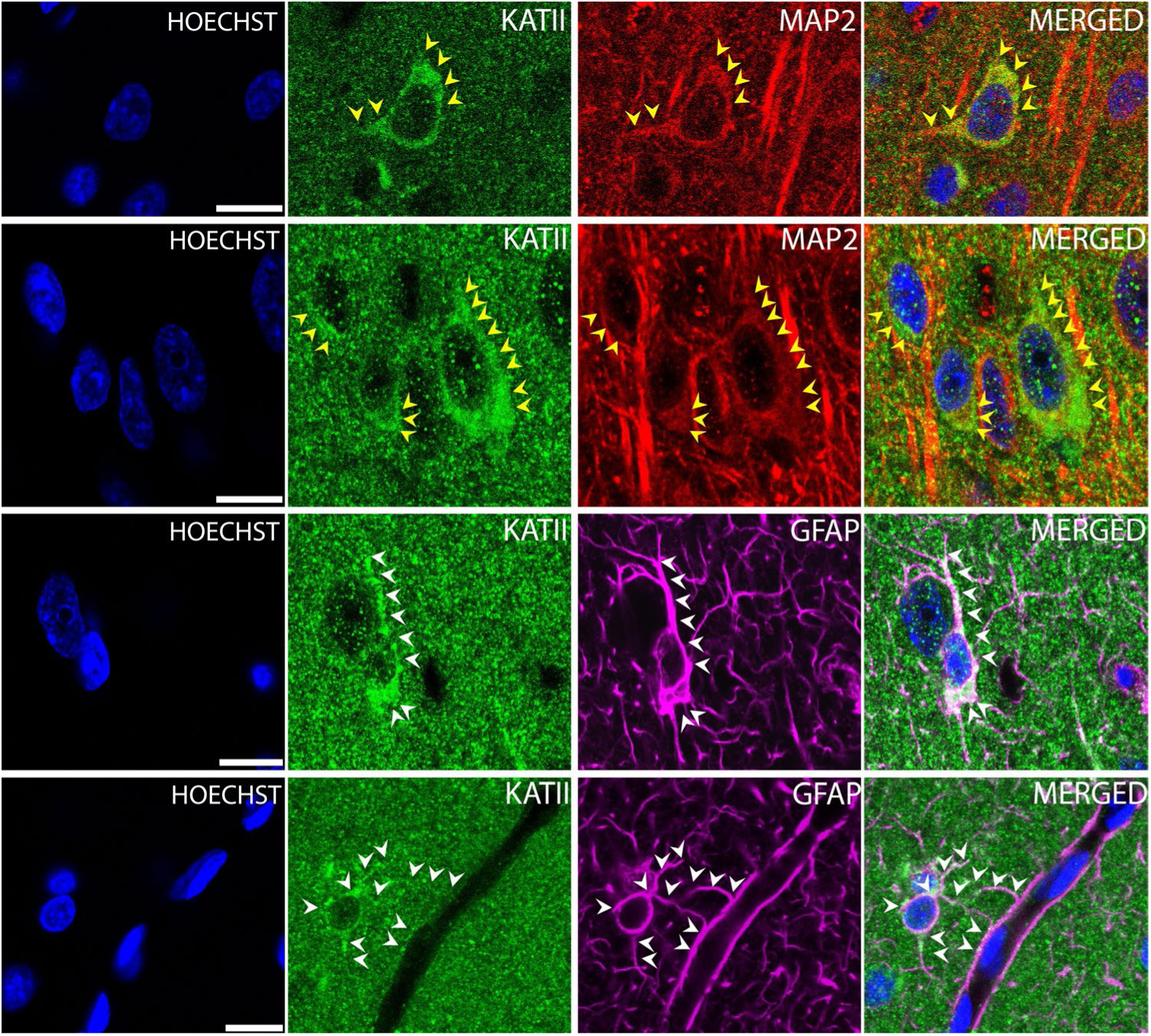
KAT II immunolabeling in layer III of aged macaque dlPFC using multiple label immunofluorescence. KAT II labeling (green) in pyramidal neurons (MAP2, red) in aged rhesus macaque dlPFC layer III. KAT II labeling is visualized in perisomatic and dendritic processes of pyramidal neurons, shown with yellow arrowheads (**a-b**). KAT II is frequently observed in the processes of astrocytes, labeled with GFAP (magenta), and outlined by the white arrowheads (**c-d**). Hoechst label (blue) indicates nuclei. Scale bars: 10μm.

Cellular and subcellular localization of KYNA-KYNA is also expressed in pyramidal cells and glia in aged macaque dlPFC (Fig. S2), and thus a more in-depth characterization was performed using immunoEM. Nanoscale analysis showed robust expression in astrocytes (Figs. 3a-g, Fig. S3b-f), especially in the astrocytic processes next to synapses on spines (peripheral astrocytic processes, i.e. PAPs), where astrocytes regulate glutamate neurotransmission. The focal expression of glial KYNA next to the synapse suggests that KYNA could be released into the synaptic cleft to interact with NMDAR and nic-α7R in the PSD (e.g. schematized in Fig. 3e). KYNA was also seen within a subset of dendritic spines (Figs. 3h-i, Fig. S3a) and pyramidal cell shafts (Fig. 3j), including on dendritic microtubules where it may be trafficking within the neuron (Fig. 3j). The expression of KYNA in spines was of particular interest, as there was often localization near the PSD in vesicular-like structures (Figs. 3h-i), suggesting that neurons may release KYNA into the synaptic cleft to interact with their own NMDAR and nic-α7R to produce auto-inhibition.

**Figure 3.**
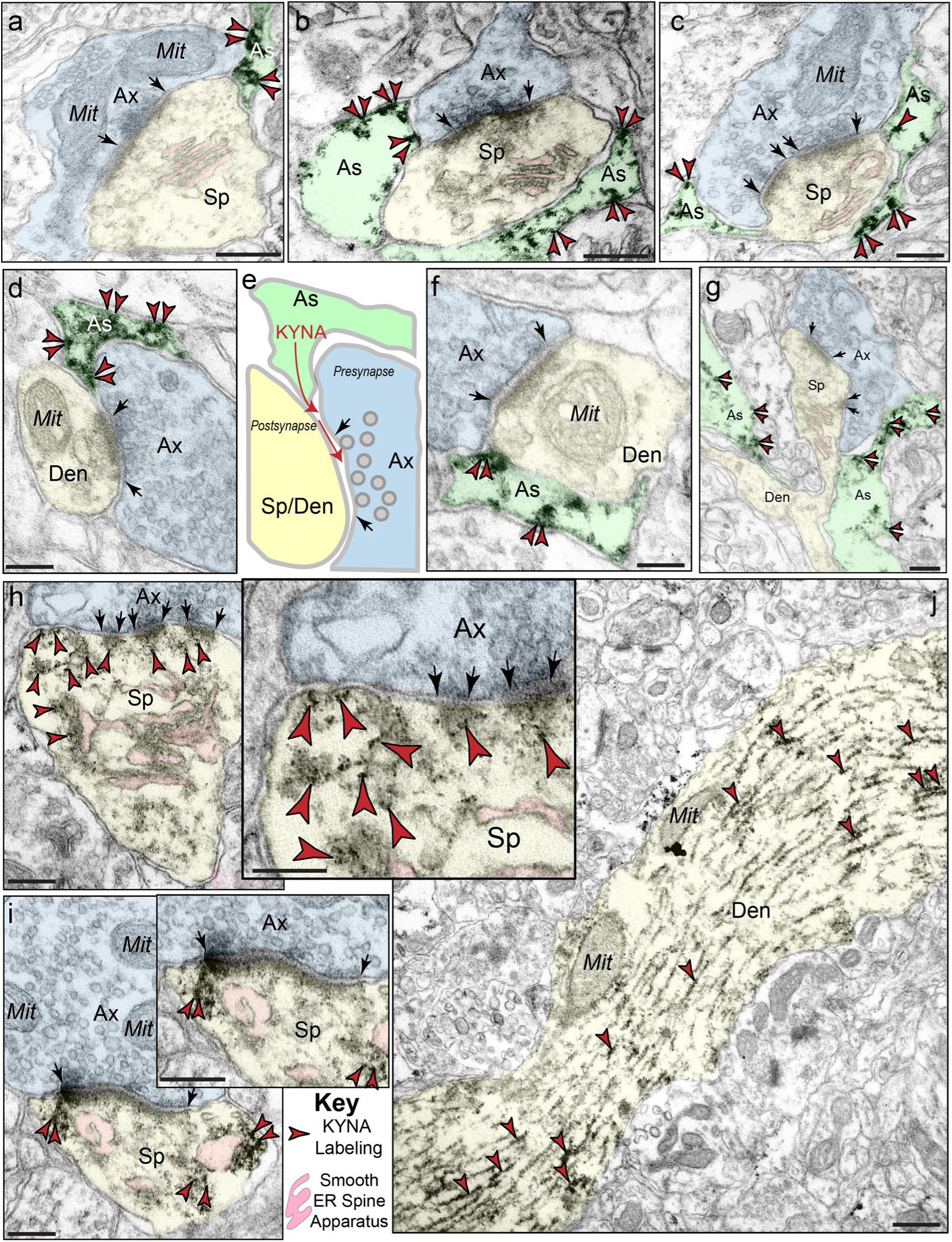
Subcellular localization of kynurenic acid in aged rhesus macaque dlPFC layer III. Kynurenic acid immunolabeling (indicated by red arrowheads) is visualized in astrocytic leaflets (pseudocolored green) ensheathing axospinous (**a-c, g**) and axodendritic (**d, f, g**), asymmetric glutamatergic synapses. Panel **e** highlights a cartoon depicting release of kynurenic acid from astroglial leaflets to engage with glutamatergic excitatory synapses in dendritic spines and dendritic shafts. Kynurenic acid immunolabeling appears to show a preference for the perisynaptic astroglial processes (PAPs) near the plasma membrane close to the excitatory glutamatergic synapse. In **g**, a dendritic spine is seen emanating from the dendritic shaft of a pyramidal cell; kynurenic acid immunolabeling is visualized in astroglial leaflets near the dendritic spine head and the shaft. Kynurenic acid immunolabeling is prominently expressed in *postsynaptic* compartments in dendritic spines (pseudocolored yellow), near the plasma membrane and PSD synaptic active zone (**h-i**). Intriguingly, kynurenic acid immunolabeling is visualized in vesicular-like structures (insets with higher magnification) within the dendritic spine head, positioned to undergo endocytosis or exocytosis. All dendritic spines receive axospinous Type I asymmetric glutamatergic-like synapses, and the spine apparatus is pseudocolored in pink in the spine head. Synapses are between arrows. In aged macaque dlPFC layer III, kynurenic acid immunolabeling was visualized in dendritic shafts (pseudocolored yellow) and was associated with microtubules oriented in parallel bundles (**j**). Kynurenic acid immunolabeling within the dendritic shaft was oftentimes in close proximity to mitochondria. Ax, axon; As, astroglial; Mit, mitochondria; Sp, dendritic spine; Den, dendrite. Scale bars, 200 nm.

### Plasma levels of kynurenine increase with age in macaques

The kynurenine/tryptophan ratio in plasma showed a significant correlation with age (r=0.63, n=15 monkeys from 11-29 years, p<0.01), increasing from 0.021 in an 11 year old monkey to 0.053 in a 29 year old monkey.

### Iontophoresis of KYNA reduces the firing of dlPFC Delay cells needed for working memory

The effects of KYNA on dlPFC physiological function were examined in a young adult (age 10-11 years, male, plasma kynurenine/tryptophan=0.021), middle-aged (age 15 years, male, plasma kynurenine/tryptophan=0.023) and aged (age 24-26 years, female, plasma kynurenine/tryptophan=0.04) macaque performing an oculomotor visuospatial working memory task (Fig. 4a). Single unit recordings were made from the dlPFC subregion most needed for task performance (Fig. 4b) and were coupled with iontophoretic administration of minute amounts of drug to alter the local neurochemical environment (Fig. 4c). Delay cells exhibit elevated and sustained neuronal firing across the delay period for their preferred direction, but not other directions (Fig. 4d), and thus are spatially tuned as defined by d’ signal detection analyses. There is a known age-related decrease in dlPFC Delay cell firing^44^, which may be related in part to endogenous KYNA expression, while exogenous application of KYNA may have greater effect in younger monkeys with their higher baseline. Indeed, iontophoresis of KYNA onto dlPFC Delay cells produced a significant, dose-related reduction in Delay cell firing in all monkeys, with the greatest effects in the younger animal, but still significant effects in the oldest animal (Fig. 4e-i; Fig.S4; Fig. S5). An example of KYNA’s effects on a Delay cell from the aged monkey are shown in Fig. 4e, where application of KYNA at a dose of 15nA had little effect on delay-related firing (p=0.99), while subsequent application of KYNA at 30nA significantly reduced delay firing for the neuron’s preferred direction only (p=0.046), reducing the spatial tuning of this delay cell (d’: control=1.01 and KYNA@30nA=0.58). Delay cell firing was restored toward control levels when KYNA was no longer applied. Consistent results were observed for the entire population of delay cells tested from the aged monkey (Fig. 4f), where KYNA (30-50nA) significantly reduced the delay firing of all 19 aged delay cells recorded. Although KYNA significantly decreased neuronal firing across the delay epoch for both the preferred and non-preferred directions, it had much greater effects for the preferred direction (2-way ANOVA-R, F(1,18)=42.22, p<0.0001), thus significantly reducing the d’ measure of spatial tuning during the delay epoch (paired t test, t=7.475, p<0.0001). In contrast, there was no direction-specific effect of KYNA on either the fixation-(Fig. 4g, p=0.3412), cue-(p=0.1564) or response-related firing (p=0.9365). A similar but even more striking effect of KYNA was seen in in the young adult monkey (Fig. 4h, firing rate; 2-way ANOVA-R, F (1, 28) = 23.72, p<0.0001; d’; paired t test, t=6.691, p<0.0001), where even very low doses (10-20nA) reduced Delay cell firing (Fig. 4i; p<0.0001), a dose range that had no effect on Delay cells from the aged monkey (p=0.384). This pattern of response is consistent with endogenous KYNA expression in the aged monkey already contributing to basal reductions in cell firing, a hypothesis supported by the experiments with the KAT II inhibitor.

**Figure 4.**
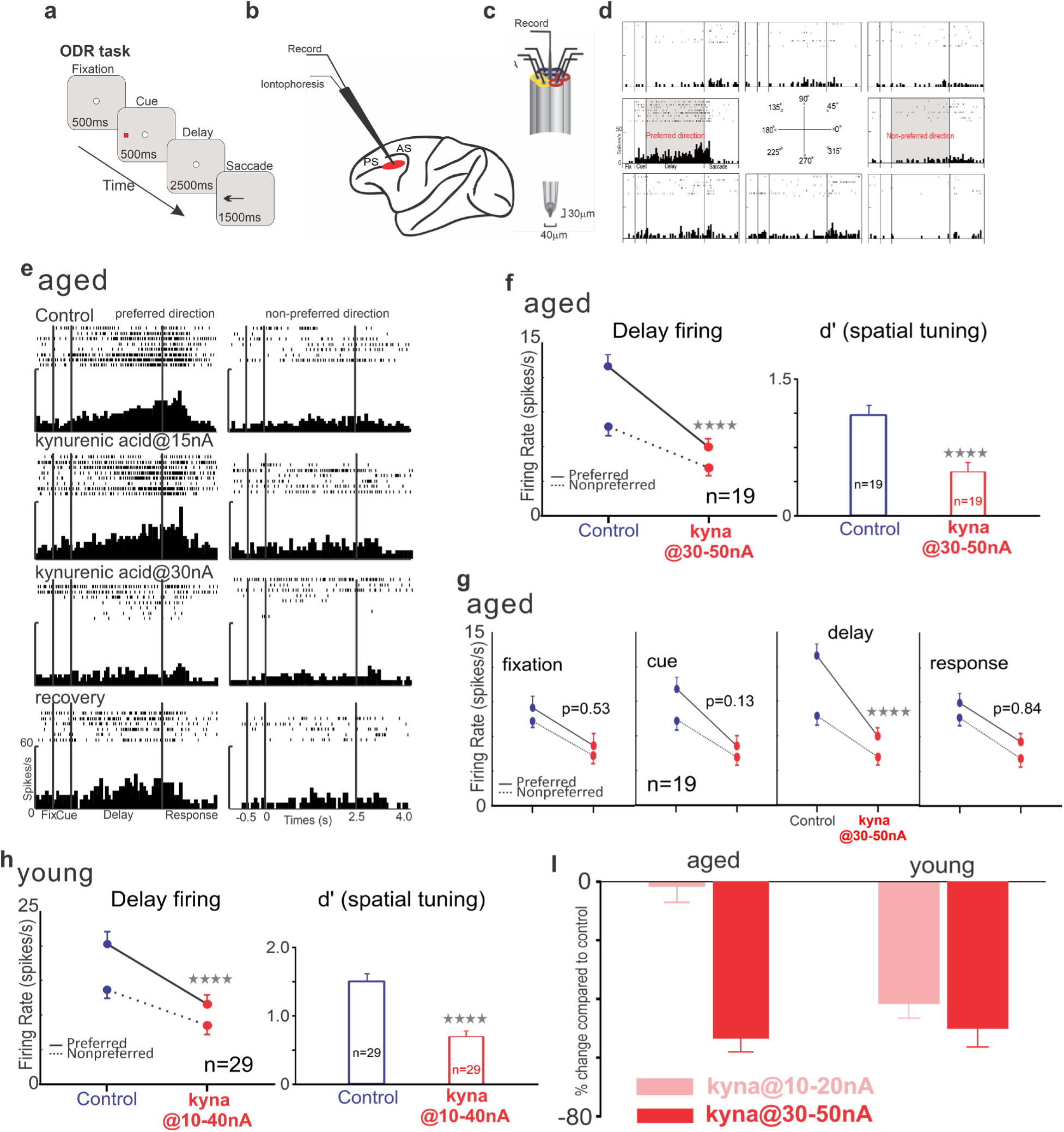
KYNA reduced delay firing and spatial tuning of dlPFC Delay cells in an aged and a young macaque. **a,** The oculomotor delayed response (ODR) task. **b,** recording locations at the posterior end of the principal sulcus in the dlPFC. **c,** iontophoretic electrode. **d,** an example of Delay cell. **e,** KYNA produced a dose-dependent reduction in firing of a Delay cell from an aged macaque. **f,** KYNA reduced the delay firing and spatial tuning of Delay cells (n=19) from an aged monkey. **g,** Further analyses showed that KYNA selectively reduced firing during the delay epoch compared to the other task epochs (n=19). **h,** KYNA markedly reduced delay firing and spatial tuning of dlPFC Delay cells in the young monkey (n=29). **i,** KYNA at higher doses (30-50nA) significantly reduced the delay-related firing of dlPFC neurons from either the young or aged monkey, while KYNA at lower doses (10-20nA) only reduced delay-related firing in the young monkey, not the aged monkey.

### Iontophoresis of KAT-II inhibitor enhanced delay firing and spatial tuning of dlPFC Delay cells with greatest effects in the aged monkey

We next examined the effect of the KAT-II inhibitor, PF-04859989 (PF) to see if it would enhance delay-related firing. PF had the converse profile, having its greatest effects in the aged monkey compared to the young and middle-aged animals (Fig. 5). Single neuron examples following iontophoresis of 30nA of PF are shown for the aged monkey (Fig. 5a) and the young monkey (Fig. 5b). Iontophoresis of PF at 40nA enhanced delay-related firing and spatial tuning, especially in the aged macaque who may have greater levels of endogenous KYNA (Fig. 5c). More subtle enhancing effects were seen with the middle-aged (Fig. 5d) and young (Fig. 5e) monkey, with the enhanced delay firing in the aged monkey significantly greater than that in the young and middle age monkeys (Fig. 5f; p<0.0001). Similar age-related effects of PF on working memory performance were seen with systemic administration (see below).

**Figure 5.**
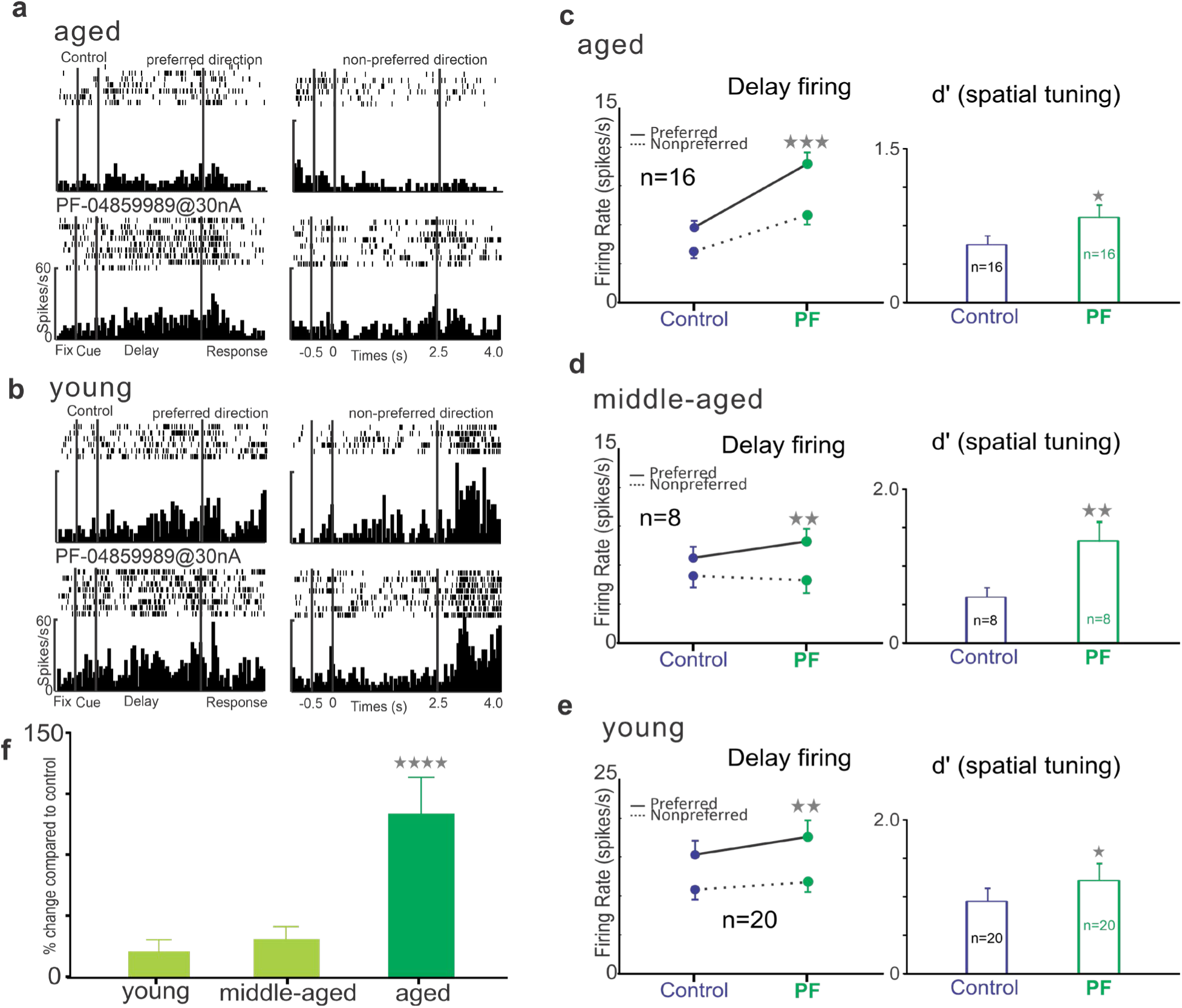
The KAT-II inhibitor PF-04859989 (PF), enhanced delay firing and spatial tuning of dlPFC Delay cells in the aged, middle-aged and young macaque, with greatest effects in the aged monkey. **a**, An example of a dlPFC Delay cell from an aged macaque showing that PF (30nA) enhanced delay-related firing. **b**, An example of a dlPFC Delay cell from a young macaque showing that PF (30nA) had only subtle effects on delay-related firing. **c**, In recordings of Delay cells from the aged dlPFC, PF (30-40nA) enhanced delay firing and spatial tuning (n=16 neurons). **d,** In recordings of Delay cells from the middle-aged dlPFC, PF (30-40nA) enhanced delay firing and spatial tuning (n=8 neurons). **e,** In recordings of Delay cells from the young dlPFC, PF (30-40nA) enhanced delay firing and spatial tuning (n=20 neurons). **f,** PF produced a significantly stronger enhancement of Delay cell firing in the aged monkey than in the young animals.

### Both NMDAR and nic-α7R blockade contribute to KYNA’s detrimental actions in primate dlPFC

There is longstanding evidence that KYNA blocks the glycine site on NMDAR, which is needed for effective glutamate binding. Thus, KYNA could reduce endogenous NMDAR transmission in dlPFC by blocking this site. D-serine is an agonist at the glycine site; thus we tested whether KYNA’s detrimental effects could blocked by co-iontophoresis with d-serine. A single neuron example from the young monkey is shown in Fig. 6a, where d-serine at 20nA alone and subsequent co-application with KYNA had no clear effect on delay firing (ordinary 2-way ANOVA, F (2, 37) = 0.3975, p=0.6748; control vs d-serine vs d-serine+KYNA) and spatial tuning (d’: control 2.68; d-serine 2.27; d-serine+kyna 2.85). However, when d-serine application was terminated, KYNA alone significantly reduced delay firing (F (2, 38) = 3.452, p=0.0419; control vs KYNA vs recovery). Similar effects were evident at the population level (Fig. 6b), where co-application of d-serine with KYNA prevented the reducing effects of KYNA (2-way ANOVA-R, F (2, 36) = 0.9167, p=0.4089; control vs d-serine vs d-serine+kyna); while when d-serine was terminated, KYNA alone consistently reduced delay firing (F (1,18) = 18.56, p=0.0004; control vs KYNA). Similar effects were seen in the aged monkey (Fig. S5). These results are consistent with KYNA reducing delay firing through blocking the glycine site on the NMDAR.

**Figure 6.**
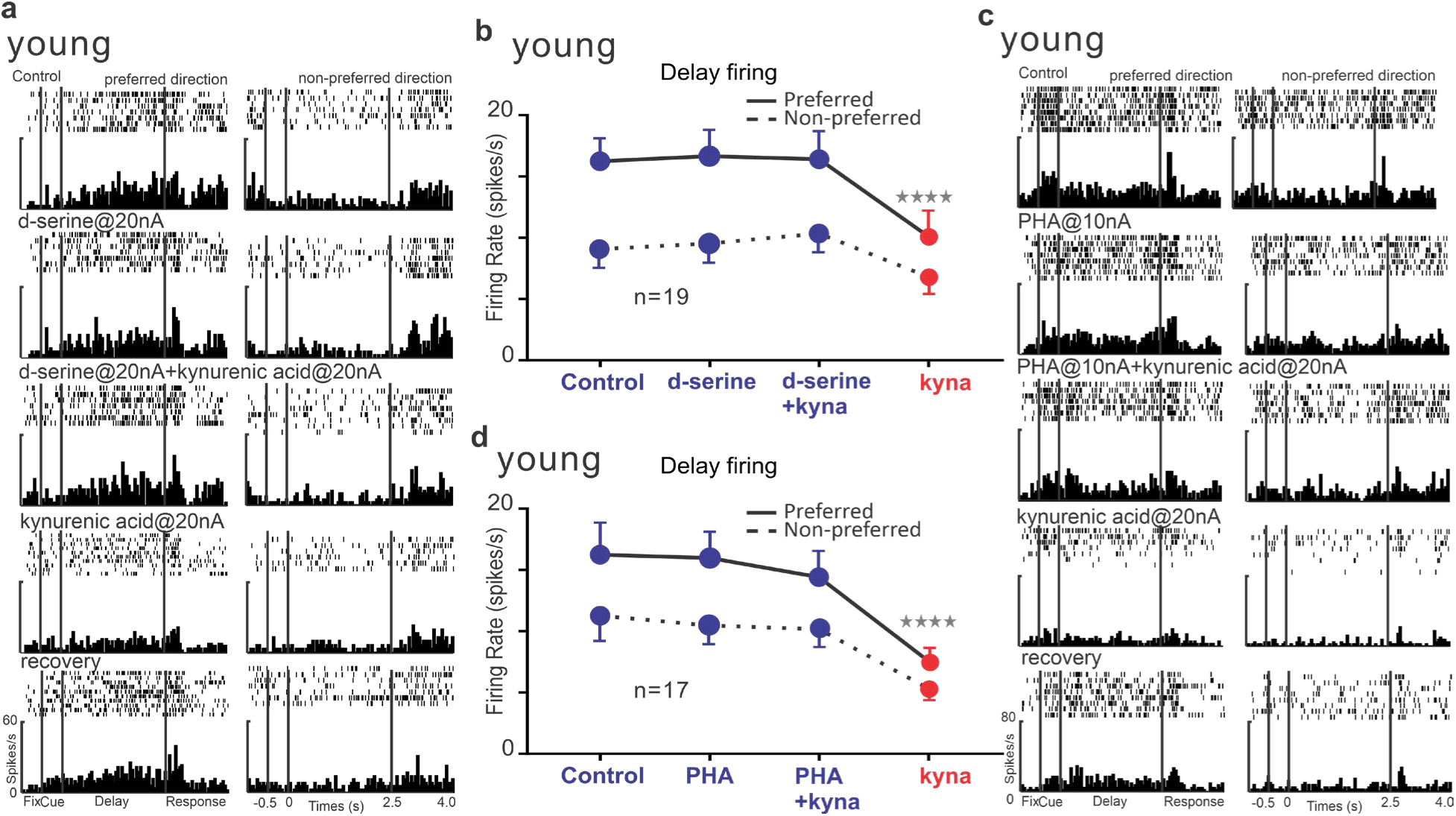
Evidence that both NMDAR and nic-a7R blockade contribute to KYNA’s detrimental actions in primate dlPFC. **a**, D-serine is an agonist at the glycine site on the NMDAR. An example neuron from a young adult monkey showing that co-iontophoresis of d-serine with KYNA prevented the reduction in delay firing that ensued when KYNA was then applied on its own. Firing began to recover when KYNA was no longer applied. **b**, The average response of all dlPFC Delay cells treated with d-serine and KYNA, where d-serine prevented the reduction in firing caused by KYNA alone (n=19 neurons). **c**, An example neuron from a young adult monkey showing that co-iontophoresis of the nic-a7R agonist, PHA, with KYNA prevented the reduction in delay firing that ensued when KYNA was then applied on its own. Firing began to recover when KYNA was no longer applied. **d**, The average response of all dlPFC Delay cells treated with PHA and KYNA, where PHA prevented the reduction in firing caused by KYNA alone (n=17 neurons).

KYNA has also been shown to block nic-α7R in some circuits, and as dlPFC Delay cell firing requires nic-α7R permissive actions for NMDAR neurotransmission, we tested for KYNA interactions at this receptor in the young monkey as well using the nic-α7R agonist, PHA 543613 (PHA). We applied a low dose (10nA) of PHA that had no effect on delay firing by itself (control vs PHA: Fig. 6c, single example, F (1, 26) = 2.264, p=0.1445; Fig. 6d, population, F (1, 16) = 0.8471, p=0.371). Co-iontophoresis of PHA with KYNA blocked KYNA’s effects on Delay cell firing and spatial tuning (PHA+kyna vs kyna: Fig. 6c, single example, F (1, 25) = 23.94, p<0.0001; Fig. 6d population, F (1, 16) = 25.24, p=0.0001), consistent with nic-α7R blockade also contributes to KYNA’s detrimental actions in primate dlPFC.

### Systemic administration of agents that reduce KYNA production improve working memory performance in aged macaques

The overarching goal of this research is to try to identify pharmacological treatments for neuroinflammatory cognitive disorders. Thus, we examined the effects of systemic administration of agents that reduce the production of KYNA on working memory performance in rhesus monkeys (actions summarized in Fig. 7f). Rhesus macaques naturally develop working memory deficits with advancing age ^46–49,53^, and we hypothesized that some of these deficits may arise from endogenous KYNA expression. Systemic administration of the KAT-II inhibitor, PF, to macaques (10-33 years) improved the performance of the aged monkeys (Figs. 7a-c), with a significant correlation between increased drug efficacy and increased age (Fig. 7a,b), and the largest effects in the oldest monkeys (Fig. 7c). These findings are consistent with the greater inflammation ^54^ and KYNA levels ^33^ in older animals.

**Figure 7.**
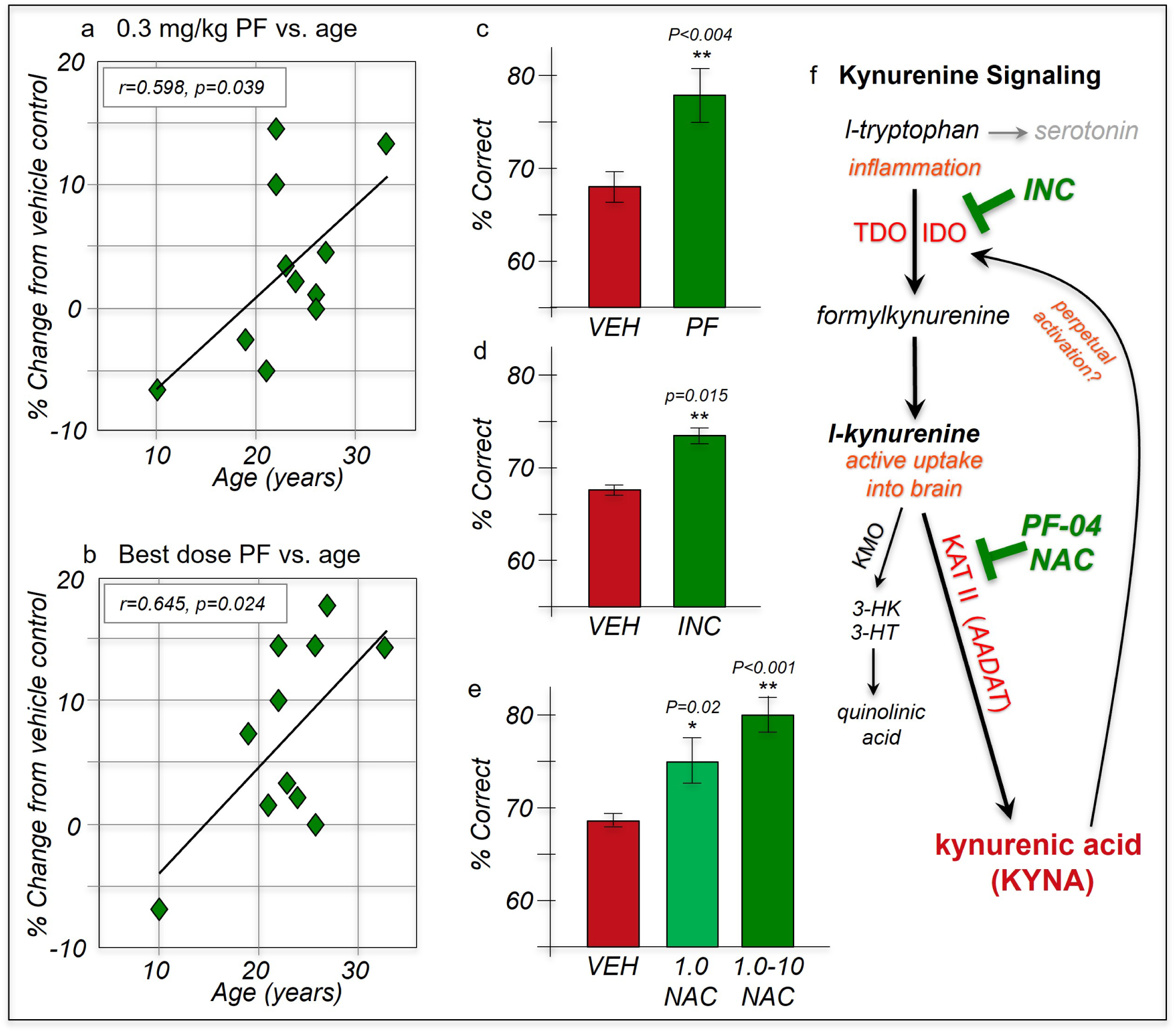
Systemic administration of agents that reduce KYNA synthesis improve working memory performance in aged macaques. **a.** The effects of the KAT-II inhibitor, PF-04859989 (PF; 0.3 mg/kg, sc, 2 hrs before testing) on working memory performance in macaques aged 10-33 years (n=11). There was a significant correlation between the degree of improvement and age, with older animals improved. **b.** The effects of the best dose of the KAT-II inhibitor, PF, between 0.003-3.0 mg/kg, (sc, 2 hrs) on working memory performance in the same animals as in panel a. There was a significant correlation between the degree of improvement and age, with older animals improved. **c.** A best dose of the KAT II inhibitor, PF (0.003-3.0 mg/kg, sc, 2 hrs), significantly improved working memory performance in aged macaques (>21 years, n=8). **d.** The IDO inhibitor, INCB024360 (0.1 mg/kg, po 2 hrs before testing) significantly improved working memory performance in aged macaques (ages 18-33 years; n=10). **e.** N-acetyl cysteine (NAC) inhibits KAT II in addition to its anti-oxidant actions and is approved for human use. NAC (1.0 mg/kg, po, 2 hrs) significantly improved working memory performance in aged macaques (20-33 years; n=10); a best dose between 1-10 mg/kg produced optimal performance for each animal.

Additional experiments focused on aged macaques and found that systemic administration of the IDO inhibitor, INCB 024360 similarly improved working memory performance compared to vehicle control (Fig. 7d). Finally, we tested N-acetylcysteine (NAC), as this agent is already approved for human use, and has recently been shown to inhibit KAT II actions (Fig. 7f) and reduce KYNA levels, in addition to its anti-oxidant properties^55,56^. An acute dose of NAC between 1-10mg/kg improved working memory performance in aged macaques, e.g. with significant improvement at 1 mg/kg compared to vehicle control (Fig. 7e). Testing of their plasma kynurenine/tryptophan levels showed elevated kynurenine (mean ratio of 0.035±0.003).

## DISCUSSION

### Summary

The current study found a large expansion of KAT II/KYNA signaling in the primate dlPFC, with expression in both neurons and glia, where KYNA produced a marked loss of neuronal firing needed for working memory and higher cognition. The loss of firing arose from KYNA blocking both NMDAR and nic-α7R, the receptors essential to dlPFC neurotransmission^23,24^. Systemic administration of agents that inhibited KAT II or kynurenine synthesis improved working memory in aged macaques with naturally occurring KYNA expression, encouraging the development of IDO or KAT II inhibitors for the treatment of inflammatory cognitive disorders such as long-COVID and schizophrenia.

An inherent weakness of i*n vivo* recording is that neuronal identity cannot be determined e.g. to distinguish pyramidal cells from nonfast-spiking interneurons. However, the majority of neurons in the dlPFC are pyramidal cells, and thus many of the recorded neurons were likely pyramidal cells. The current recording methods also cannot determine laminar identity, but previous studies have indicated that Delay cells are mostly found in superficial layers^57^, and layer III is particularly large in macaque and human dlPFC^58,59^.

### Species differences- expansion of kynurenine KAT II signaling in primates

The current study found striking species differences in the expression of *AADAT* encoding the synthetic enzyme for KYNA, KAT II. In general, there was very low expression in mouse frontal cortex, with expression focused in glia. In contrast, there was much higher expression in macaques and humans, with the largest levels in a subgroup of pyramidal cells, followed by high levels in astrocytes and oligodendrocytes. The current data from primates are in line with a previous analysis of astrocytes showing that *AADAT* expression is increased almost 80-fold in humans compared to mice^60^, and that KYNA levels in PFC are 10-20 times greater in humans than rats^61^. The current transcriptomic data from mice are also consistent with previous studies of protein expression in mPFC of rodent, where IHC studies of KAT II showed focal localization in glia^62,63^, as well as some interneurons^63^. In contrast, the transcriptomics found prominent *AADAT* expression in both neurons and glia in macaque and human dlPFC, with excitatory neurons having the highest levels, and prominent protein expression of KAT II and KYNA in both neurons and glia in aged macaque dlPFC. These species differences in neuronal *AADAT* expression -mostly interneurons in mice, mostly excitatory cells in human and nonhuman primates-suggests that the overall effects of KYNA inflammation would differ markedly between species, with a blockade of NMDAR excitation of GABA interneurons in mice leading to overexcitation of circuits, and a predominant blockade of NMDAR excitation of pyramidal cells in primates leading to underexcitation of circuits^19^. The predominance of KYNA actions on pyramidal cell dendritic spines was reinforced by the immunoEM of macaque dlPFC, showing prominent expression of KYNA near excitatory synapses on spines, positioned to disrupt the NMDAR and nic-a7R^23,24^ neurotransmission that mediates working memory, confirmed by the physiological data showing marked loss of dlPFC neuronal firing with local KYNA application. The great expansion and qualitative differences in KAT II kynurenic acid signaling in primate cortex, emphasize why it is so important to have primate models of inflammation, especially if we are to understand and treat higher cognitive deficits in disorders such as long-COVID, schizophrenia and AD that have an inflammatory etiology.

The large species differences in the capacity to synthesize KYNA are intriguing, given that KYNA blocks NMDAR, and the expression of NMDAR-GluN2B (*GRIN2B*) also expands in primate dlPFC^64^, increasing across the cortical hierarchy^65,66^. For example, mPFC neurons in rodent have a large dependence on AMPAR as well as NMDAR neurotransmission^67^, and systemic kynurenine produces only a subtle impairment in the working memory of rats^68^. Thus, KYNA inflammatory signaling in primates may have expanded in concert with increased NMDAR actions to ensure that dlPFC was silenced under inflammatory conditions when energy is needed elsewhere. The current data also emphasize why mouse models can be inadequate for accurately capturing the etiological roles of inflammation in human cognitive disorders, and why the role of KYNA in inflammatory signaling sometimes may be underestimated if not performed in primate dlPFC neurons.

The expansion of KAT II KYNA signaling in primates may also have great relevance to perinatal inflammatory insults and the development of cortical circuits, as NMDAR are required for the creation of appropriate cortical connections^69^. Thus, expansive KYNA blockade of these receptors while cortical connections are forming may produce wide-ranging deficits in connectivity that could contribute to intellectual disability and other mental disorders, which may be more subtle in rodent models^70^.

### Potential roles of KYNA in neurons

The presence of KYNA and *AADAT* in pyramidal cells in macaque dlPFC was unexpected, given their focused localization in glia in rodent cortex^62^. The role of intraneuronal KYNA in pyramidal cells is unknown and an area for future research. Presumably, KYNA is only effective in blocking NMDAR and nic-a7R neurotransmission when it is extracellular, e.g. within the synaptic cleft. The immunoEM indicates that KYNA may be localized within vesicular-like structures near the PSD (e.g. Fig. 3g), suggesting that it might be released into the synaptic cleft to reduce NMDAR/nic-a7R neurotransmission under inflammatory conditions, thus producing “auto-inhibition”. Conversely, these may be sites of KYNA uptake from the synapse, suggesting a more dynamic relationship.

The transcriptomic analyses of macaque dlPFC showed that the very highest levels of *AADAT* were in a subset of pyramidal cells that co-expressed *RORB*. Although RORB is a marker of layer IV neurons in mouse cortex^71^, it is expressed in pyramidal cells across multiple layers in macaque dlPFC, including those in layer III (Fig. S1). These data suggest that KAT II can be synthesized within neurons throughout all layers of macaque dlPFC, which may provide widespread reduction in NMDAR-dependent excitatory actions. As interneurons in adult primate and rodent PFC rely more on AMPAR than NMDAR^72–74^, KYNA expression may be especially detrimental to the firing of pyramidal cells.

### A new view on kynurenic acid as the “protective” metabolite

Traditionally, kynurenine signaling has been examined within the context of the “excitotoxicity” that can occur with very high levels of glutamate release, and where increased kynurenine metabolism to QUIN further drives NMDAR stimulation, calcium entry and overload of mitochondria, and subsequent apoptotic cell death e.g.^40^. Under these conditions, KYNA protects neurons from overexcitation and apoptosis by blocking calcium entry through NMDAR^40^. This mechanism is thought to mediate the neuronal death that occurs during a stroke, or other severe, acute injuries to the nervous system^2,75^. Modeling of these toxic events is usually performed in rodent models, or in cell cultures made from rodent neurons e.g.^76^, where healthy neurotransmission has a major AMPAR component e.g.^77^.

The current study shows that in the recently evolved dlPFC circuits that rely heavily on NMDAR and nic-α7R neurotransmission for healthy function, KYNA’s blockade of NMDAR and nic-α7R is not always protective, but instead, can contribute to loss of firing and impaired cognition. This would be particularly true under conditions of normal or reduced glutamate release, as is likely the case with aging^44^, and in long-term inflammatory disorders where pathology occurs more gradually, e.g. with loss of neuronal firing, atrophy, and slow, autophagic degeneration rather than apoptosis. As these conditions are common in human age-related and inflammatory disorders including AD^78^, the current data caution that a revised, more complex view of KYNA actions is needed to understand and treat cognitive deficits in humans.

### Relevance to cognitive deficits in long-COVID (PASC)

The cognitive deficits of long-COVID consistently target dlPFC functioning^11–16^, and emerging brain imaging studies of patients engaged in cognitive tasks find alterations in dlPFC activity^79,80^. Patients who died from COVID have increased kynurenine signaling ^81^, including increased KYNA in brain^34^. Fluid biomarker data from patients with long-COVID are just beginning to emerge, but recent findings show that plasma kynurenine levels correlate with symptoms of cognitive deficits^5^, as well as depression and anxiety^6,82^, the latter which may be related to decreased dlPFC top-down control of emotion^83^. Interestingly, KYNA can perpetuate kynurenine signaling by activating IDO metabolism of tryptophan^81^, as schematically illustrated in Figure 7f, and this may sustain symptoms even after the initial cytokine inflammatory response is over^84^. The current data show that the increased plasma kynurenine levels consistently found in patients with long-COVID^4–6,81,85,86^ could directly impair higher cognitive functioning and top-down regulation of emotion through active kynurenine uptake into the dlPFC and its metabolism to KYNA, reducing dlPFC neuronal firing needed for cognition and top-down control. This interpretation would be consonant with the report by Cysique et al, where cognitive deficits in long-COVID correlated best with levels of plasma kynurenine^5^.

### Relevance to age-related cognitive dysfunction and AD

Longstanding data from rodents, macaques and humans have shown increased KYNA levels in CNS with advanced age^31–33,85^, consistent with greater inflammation with age^9,86,87^. The current data are consonant with these findings, as aged macaques were more responsive to KAT-II inhibitors than younger adult macaques, and showed greater levels of plasma kynurenine. The improvement in both Delay cell firing and working memory performance in aged macaques treated with KAT-II inhibitors suggests that endogenous KYNA contributes to cognitive deficits with age. The improved working memory with PF in the current study is coherent with previous data showing that this compound partially protected against working memory deficits induced by the NMDAR antagonist, ketamine, suggesting competitive receptor interactions^88^. It is noteworthy that the loss of dlPFC Delay cell firing with advanced age also involves dysregulated calcium-cAMP signaling, increasing the open state of potassium channels on spines^44,89,90^. Thus, calcium dysregulation can occur within the context of reduced, not increased, neuronal firing, and involves dysregulation of internal calcium release^90^, emphasizing the important qualitative differences between the aging process in dlPFC circuits and the studies of excitotoxicity commonly performed in cell cultures.

KYNA is also increased in early stages of Alzheimer’s disease^38,91^, where increased kynurenine and its metabolites in plasma correlate with measures of Aβ and neurofilament light chain assays of degeneration^92^. IDO expression is also increased near plaques and tangles in AD brain samples^37^, consistent with inflammatory contributions to sporadic AD as well as other neurodegenerative disorders^93^. In this regard, it is of interest that patients who died of severe COVID infection exhibited high levels of phosphorylated tau as well as elevated KYNA in their brains^34^.

### Relevance to schizophrenia

Schizophrenia is increasingly linked to increased inflammatory signaling e.g.^94–98^ which may contribute to the decreased dlPFC dendrites and spines^99^. Postmortem brain analyses have shown elevated KYNA levels^8,35,61^, and increased message for TDO and KAT II, in the dlPFC of patients with schizophrenia, especially in those with high cytokine levels^8^, Studies of living patients found elevated kynurenine in the plasma of those with high cytokines which correlated with impaired attentional regulation and reduced dlPFC volume^8^.

Taken all together, these findings suggest that agents that inhibit the production of kynurenine and/or KYNA would be helpful in treating the cognitive deficits of long-COVID. Currently, there are no selective TDO, IDO or KAT-II inhibitors approved for human use. However, recent data show that NAC, in addition to its anti-oxidant properties, inhibits KAT-II^55,56^. NAC is FDA-approved for treating acetaminophen overdose, and is being tested for potential protective effects in schizophrenia where it may improve working memory^100^, especially with long-term use^101^, and open-label evaluations treating longstanding traumatic brain injury^102^ or long-COVID^103^ with encouraging results. The current data suggest that more targeted compounds may be even more effective, especially given the expansion of the KYNA inflammatory pathway in the primate dlPFC.

## ACKNOWLEDGEMENTS

We would like to thank Lisa Ciavarella, Tracy White Sadlon, Sam Johnson and Michelle Wilson for their invaluable technical expertise. This work was supported by National Institute of Aging RF1 AG083090 to MW and National Science Foundation grants 2015276 to AFTA.

## AUTHOR CONTRIBUTIONS

Conceptualization, A.A. and M.W.

Methodology, S.Y., D.D., F.K., E.L., E.W., A.M., G.A., V.G. and M.W.

Investigation, S.Y., D.D., F.K., E.L., E.W., A.M., G.A., V.G. and M.W.

Writing – Original Draft, A.A. and M.W.

Writing – Review & Editing, S.Y., D.D., F.K., E.L., E.W., A.M., G.A., V.G., G.G., D.L., S.M., A.A. and M.W.

Funding Acquisition, A.A. and M.W. Resources, A.A. and M.W.

Supervision, G.G., D.L., S.M., A.A. and M.W.

## STAR Methods

### LEAD CONTACT AND MATERIALS AVAILABILITY

Further information and requests for resources and reagents should be directed to and will be fulfilled by the Lead Contact, Min Wang.

This study did not generate any new unique reagents.

### EXPERIMENTAL MODEL AND SUBJECT DETAILS

The electrophysiology experiments were performed with one young (age 10, male), one middle-aged (age 15, male) and one aged (age 24, female) Rhesus macaque. The systemic behavioral experiments were performed with 21 (10-33 years) Rhesus macaques (9 male, 12 female). Subjects were individually or pair-housed with 12-hour light cycles in humidity and temperature-controlled environments. The immunoelectron microscopy experiments were performed with three aged Rhesus macaques (age 24, 30 and 31, female). Immunofluorescence staining was carried out on free-floating sections in two aged rhesus macaques (age 28, male; age 30 female). All research was conducted according to USDA and NIH guidelines and approved by the Yale or University of Pittsburgh IACUC (for macaques), the Princeton University IACUC (for mice), and the Partners Human Research Committee (for humans). For human dlPFC tissues, brain donors were recruited by the Harvard Brain Tissue Resource Center/NIH NeuroBioBank (HBTRC/NBB), in a community-based manner, across the USA. Human brain tissue was obtained from the HBTRC/NBB. The HBTRC procedures for informed consent by the donor’s legal next-of-kin and distribution of de-identified post-mortem tissue samples and demographic and clinical data for research purposes are approved by the Mass General Brigham Institutional Review Board. Post-mortem tissue collection followed the provisions of the United States Uniform Anatomical Gift Act of 2006 described in the California Health and Safety Code section 7150 and other applicable state and federal laws and regulations. Federal regulation 45 CFR 46 and associated guidance indicates that the generation of data from de-identified post-mortem specimens does not constitute human participant research that requires institutional review board review. The details of macaques used are described in each section below; all macaques were housed under standard laboratory conditions with daily veterinary evaluation and daily enrichment including fresh fruits and vegetables and optimal paired housing.

### METHOD DETAILS

#### Transcriptomics

We obtained *AADAT* expression from the following previously published single cell/nucleus RNA sequencing datasets: mouse frontal cortex^50^; dlPFC (BA46) of neurotypical postmortem human donors^52^; and dlPFC (BA46) of macaque monkeys^51^. For each dataset, preprocessing and clustering assignments were retained as in the original publications. In Ling et al^52^, human clusters were matched to a previous human snRNA-seq dataset of middle temporal gyrus^104^ using scPred (https://github.com/powellgenomicslab/scPred). Macaque and mouse clusters were matched to human based on known marker genes (e.g. *PVALB* and *GAD1* expression for *PVALB*+ interneurons) as well as the mouse-human mapping shown in Hodge et al. When subtype clusters were present in the macaque or mouse (e.g. multiple *PVALB*+ subclusters), gene expression values were averaged across subtype. For cell type labels in Fig. 1, we adopted the approach to cross-species naming convention in Hodge et al of including putative projection type for excitatory neurons as inferred from projection-mapping studies in mouse (see Fig. 5e^104^). The layer distribution label annotation is inferred from human ISH^104^. To produce estimates of *AADAT* expression for each species (Fig 1), transcripts for all genes and for each biological replicate were first summed across cells contained in each cell type. The resulting “metacell” counts were normalized to the total number of transcripts, then scaled to counts per 100,000 transformed. Mouse data are available at dropviz.org. Human data can be found in NeMO under accession number nemo:dat-bmx7s1t^52^.

#### Immunohistochemistry

##### Animals and Tissue Preparation

Tissues from our brain bank of three aged (24, 30 and 31) rhesus macaques (*Macaca mulatta*) were used for this study. As described previously ^105–108^, rhesus macaques were deeply anesthetized prior to transcardial perfusion of 100 mM phosphate-buffer saline (PBS), followed by 4% paraformaldehyde/0.05% glutaraldehyde in 100 mM PBS. Following perfusion, a craniotomy was performed, and the entire brain was removed and dissected, including a frontal block containing the primary region of interest. The brains were sectioned coronally at 60 μm on a vibratome (Leica) across the entire rostrocaudal extent of the dorsolateral prefrontal cortex (dlPFC; Walker’s area 46). The sections were cryoprotected through increasing concentrations of sucrose solution (10%, 20% and 30% each for overnight), cooled rapidly using liquid nitrogen and stored at −80°C. Sections of dlPFC were processed for immunocytochemistry. To enable penetration of immunoreagents, all sections went through 3 freeze-thaw cycles in liquid nitrogen. Non-specific reactivity was suppressed with 10% normal goat serum (NGS) and 5% bovine serum albumin (BSA), and antibody penetration was enhanced with 0.3% Triton X-100 in 50 mM Tris-buffered saline (TBS).

##### Histology and Immunoreagents

We used previously well-characterized primary polyclonal anti-kynurenic acid antibody (1:100; ab37105; Abcam, Cambridge, MA, USA) raised in rabbit. The antibody is specific and has been validated using a range of applications, including immunohistochemistry. For example, previous studies have used the anti-kynurenic acid antibody has been used in immunohistochemistry studies in stratum pyramidale CA1 region of the hippocampus to evaluate excitotoxic mechanisms after transient forebrain ischemia. The anti-kynurenic acid antibody has also been used in the rat brain to evaluate the redox properties of L-KYN with scavenging activity assays.

##### Single-label immunoperoxidase immunohistochemistry

For single-label immunoperoxidase immunohistochemistry, sections of dlPFC were transferred for 1 hr to Tris-buffered saline (TBS) containing 5% bovine serum albumin, plus 0.05% Triton X-100 to block non-specific reactivity, and incubated in primary antibody in TBS for 72 hr at 4°C. The tissue sections were incubated in goat anti-rabbit biotinylated antibody (Vector Laboratories) at 1:300 in TBS for 2 hr, and developed using the Elite ABC kit (Vector Laboratories) and diaminobenzidine (DAB) as a chromogen. Omission of the primary antibody eliminated all labeling. Sections were mounted on microscope slides and dlPFC cortical layers were photographed under an Olympus BX51 microscope equipped with a Zeiss AxioCam CCD camera. Zeiss AxioVision imaging software was used for imaging and data acquisition.

##### Single pre-embedding peroxidase immunocytochemistry

As described previously^105^, the sections were incubated for 72 h at 4 °C with primary antibodies in TBS, and transferred for 2 h at room temperature to species-specific biotinylated Fab’ or F(ab’)_2_ fragments in TBS. In order to reveal immunoperoxidase labeling, sections were incubated with the avidin-biotin peroxidase complex (ABC) (1:300; Vector Laboratories, Burlingame, CA, United States of America) and then visualized in 0.025% Ni-intensified 3,3-diaminobenzidine tetrahydrochloride (DAB; Sigma Aldrich, St. Louis, MO, United States of America) as a chromogen in 100mM PB with the addition of 0.005% hydrogen peroxide for 10-12 minutes. After the DAB reaction, sections were exposed to osmification (concentration 1%), dehydration through a series of increasing ethanol concentrations (70-100%) and infiltrated with propylene oxide. Tissue sections were counterstained with 1% uranyl acetate in 70% ethanol. Standard epoxy resin embedding followed typical immunoEM procedures followed by polymerization at 60°C for 60 h. Omission of primary antibodies or substitution with non-immune serum resulted in complete lack of immunoperoxidase labeling. Similarly, labeling was nullified when blocking the biotinylated probes with avidin/biotin.

##### Electron microscopy and data analysis

For single-label immunoperoxidase immunohistochemistry, sections were mounted on microscope slides dlPFC layer III was photographed under an Olympus BX51 microscope equipped with a Zeiss AxioCam CCD camera. Zeiss AxioVision imaging software was used for imaging and data acquisition. All sections were processed as previously described and immunoEM imaging was conducted in dlPFC layer III ^105^. Briefly, blocks containing dlPFC layer III were sampled and mounted onto resin blocks. The specimens were cut into 50 nm sections using an ultramicrotome (Leica) and analyzed under a Talos L120C transmission electron microscope (Thermo Fisher Scientific). Several plastic blocks of each brain were examined using the 4^th^ to 12^th^ surface-most sections of each block (i.e., 200-600 nm), in order to sample the superficial component of sections, avoiding penetration artifacts. Structures were digitally captured at x25,000-x75,000 magnification with a Ceta CMOS camera and individual panels were adjusted for brightness and contrast using Adobe Photoshop and Illustrator CC.2020.01 image editing software (Adobe Systems Inc.). Approximately, 1000 micrographs of selected areas of neuropil with immunopositive profiles were used for analyses with well-defined criteria. For profile identification, we adopted the criteria summarized in Peters et al. 1991^109^.

##### Immunofluorescence on macaque tissue

Immunofluorescence staining was carried out on free-floating sections from tissues from our brain bank of two aged rhesus macaques (male: 28yo, female: 30yo). The sections were washed in 1X TBS (pH 7.4, 4 x 15 min) for 1 h at RT. Antigen retrieval was performed using Antigen Unmasking Solution (Vector Labs, H-3300-250) in a steam bath for 25min. The tissue was cooled on the bench top for 15min, followed by several washes using de-ionized water and tap water. To reduce background autofluorescence the tissue sections were immersed in fresh 0.5% Sodium Borohydride in 1X TBS for 10min at RT. The sections were washed in 1X TBS (pH 7.4, 4 x 15 minutes) for 1 h at RT. The sections were blocked for 1 h at RT in 1X TBS containing 5% bovine serum albumin, 2% Triton X-100, and 10% normal goat serum. Sections were incubated for 72 h at 4°C with MAP2 (1:1000; Abcam, Cat # ab5392), GFAP (1:1000; NeuroMab, Cat #75-240), and KATII (1:200; Proteintech, Cat #13031-1-AP) in dilution buffer (2% bovine serum albumin, 2% Triton X-100, and 1% normal goat serum). Appropriate secondary antibodies in dilution buffer were used at 1:100 for 3 h at RT (goat *anti*-rabbit AF488, goat *anti*-chicken AF647, goat *anti*-mouse AF594). All subsequent steps were performed in the dark and at RT. The sections were first washed in 1X TBS (pH 7.4, 4 x 15 minutes) and counterstained with Hoechst 33342 for 10 minutes (1:10,000, ThermoFisher, Cat# H3570). The sections were washed in 1X TBS (pH 7.4, 3 × 10 minutes) before mounting onto slides using ProLong Gold Antifade Mountant (Invitrogen, Cat# P36930).

##### Confocal imaging and processing

Confocal images were acquired using a Leica TCS SP8 Confocal Laser Scanning Microscope (inverted), with the HC PL APO 40X/1.40 oil white objective (Leica), HC PL APO 20X/0.75 NA dry white objective (Leica), and HCX PL APO CS 63X/1.40 oil white objective (Leica). Images were obtained under laser excitation at 407nm, 488nm, 543nm, and 633nm. Emission filter bandwidths and sequential scanning acquisition were set up in order to avoid possible spectral overlap between fluorophores. The confocal Z-stacks were processed into maximum intensity Z-projections using Fiji. Images were labeled and assembled into a figure using Adobe Photoshop CS5 Extended (version 25.5.1 ×64, Adobe Systems Incorporated) and Adobe Illustrator (version 28.3, Adobe Systems Incorporated).

#### Plasma levels of kynurenine and tryptophan

Plasma levels of kynurenine and tryptophan were determined to calculate the kynurenine/tryptophan ratio from macaques who were still alive at the time of the experiment and used in the physiology and behavior experiments. Plasma tryptophan and kynurenine were assayed by high performance liquid chromatography (HPLC) using a modification of the method of Widner et al.^110^. Briefly, 50 uL plasma samples were deproteinized with 50 uL of 5% perchloric acid after addition of 5 uL of 5% ascorbic acid and 50 ng (5 uL) of the internal standard, 5-hydroxytryptophan. Supernatants were directly injected on a 15 x 0.46 cm 5 um Microsorb C18 column eluted (0.9 mL/minute) with a mobile phase composed of 96% pH 3.4, 1.5% aqueous acetic acid (pH adjusted with 1 M sodium hydroxide) and 4% methanol (added after pH adjustment). The compounds were measured fluorometrically (tryptophan, 285/345 nm excitation/ emission wavelengths, Shimadzu 10Axl) and by ultraviolet absorbance (kynurenine, 360 nm, Hitachi L-7400), with within-day and day-to-day coefficients of variation of less than 6%.

#### Electrophysiology and Iontophoresis

##### Oculomotor delayed response (ODR) task

Rhesus macaques were trained to perform an ODR task, a test of visuospatial working memory. The task requires the subject to make a memory guided saccade to a remembered visuospatial location for a juice reward. The task is illustrated in Fig.4A. A central light is illuminated on an LED display monitor, serving as a fixation target. To initiate a trial the subject maintains fixation at the central spot for 0.5s (fixation period). Following this fixation, a cue is illuminated for 0.5s (cue period) at 1 of 8 peripheral targets located at an eccentricity of 13° with respect to the fixation spot. After the cue is extinguished, a 2.5s delay period follows (delay period). The subject is required to maintain fixation on the central spot throughout both the cue presentation and the delay period. At the end of the delay period the fixation spot is extinguished, instructing the animal to make a memory-guided saccade to the remembered location. A trial was considered successful if the animal made a saccade to the area within 2° around the previously cued location within 0.5s after the offset of the fixation spot. If completed successfully the animal was rewarded with juice immediately after the successful response. The inter-trial interval was 3s. The animal’s eye position was monitored with ISCAN Eye Movement Monitoring System, and the ODR task was generated by PictoBox System (developed by Dr. Daeyeol Lee and colleagues, Yale University).

##### Recording Site

Animals underwent a magnetic resonance imaging (MRI) brain scan to obtain the exact anatomical coordinates of the desired recording site over the caudal principal sulcus of cortical area 46 shown in Fig. 4B. These coordinates were then used to guide placement of the chronic recording chambers and electrophysiological recordings.

##### Recording and Iontophoresis

Electrodes (Fig. 4c) for dual recording and iontophoresis were constructed with a 20μm-pitch carbon fiber inserted into the central barrel of a 7-barrel nonfilamented capillary glass (Friedrich and Dimmock). The assembly was pulled using a custom electrode puller (PMP-107, Microdata Instrument Inc.) and the tip was beveled to reach impedances of 0.3-1.0 MΩ with tip sizes of 30-40μm. The outer 6 barrels of the electrode were then filled with up to 3 different drug solutions (2 consecutive barrels for each drug), which were pushed through the tip of the electrode using air. A Neurophore BH2 iontophoretic system (Medical Systems Corp.) was used to deliver the drugs. Drugs were ejected at currents that varied from 10nA – 50nA. Retaining currents of 5nA at the opposite polarity were used in a cycled manner (1s ON, 1s OFF) when not applying drugs. Drug ejection did not create noise in the recording, and there was no iontophoresis-related change in spike waveforms at any ejection current. The pharmacological agents used for iontophoresis can be found in the Key Resources Table. Each compound was dissolved at 0.01 M concentration in either sterile water or saline with pH 3-4 or pH 8-9. **Our previous saline control experiments as well as the saline control experiments from other labs showed that saline with the same pH had no specific effects on firing rate or spatial tuning of delay cells.**

The electrode was mounted on a MO-95 micromanipulator (Narishige, East Meadow, NY) in a 25-guage stainless steel guide tube. *Dura mater* was punctured using the guide tube to enable access of the electrode to cortex. Extra-cellular voltage was amplified using an AC/DC differential preamplifier (Model 3000, A-M SYSTEMS) and band-pass filtered (180Hz-6Hz, 20dB gain, 4-pole Butterworth; Kron-Hite, Avon, MA). Signals were digitized (15kHz; micro 1401, Cambridge Electronics Design, Cambridge, UK) and acquired using the Spike2 software (CED, Cambridge, UK). Neuronal activity was analyzed using waveform sorting by a template-matching algorithm. Post-stimulus time histograms (PSTHs) and rasters were constructed online to determine the relationship of unit activity to the task. Unit activity was measured in spikes per second. If the rasters showed that a neuron displayed task-related activity, recording continued and pharmacological testing was performed.

Neuronal activities were first collected from the cell under a control condition in which at least eight trials at each of eight cue locations were obtained. A typical Delay cell is shown in Fig. 4D. Upon establishing the stability of the cells’ activity, this control condition was followed by iontophoretic application of drug(s). Dose-dependent effects of the drug were tested in two or more consecutive conditions, followed by a Recovery condition or a Reversal condition. Drugs were continuously applied at a relevant current throughout a given condition. Each condition had ∼8 (6-12) trials at each location to allow for statistical analyses of drug effects.

Each experiment used multiple drug application conditions and thus required the aged monkey to perform at least 300 trials for each recording. Aged monkeys perform a limited number of trials compared to young adults, and often stop working in the middle of a recording session**. A total of 132 delay cells from a total of 88 recording sessions were successfully tested with all drug treatments. Of these, 35 were from the aged macaque (age 24, female), 12 from the middle-aged macaque (age 15, male) and 85 from the young macaque (age 10, male).**

##### Electrophysiology Data Analyses

Each trial in the ODR task was divided into four epochs – initial Fixation, Cue, Delay and Response (Saccade). The initial Fixation epoch lasted for 0.5 sec. The Cue epoch lasted for 0.5 sec and corresponds to the stimulus presentation phase of the task. The Delay lasted for 2.5 sec and reflects the mnemonic component of the task. The Response phase started immediately after the Delay epoch and lasted ∼1.5 sec. Data analysis was performed in MATLAB, SPSS and GraphPad Prism 7.01. This study focused on Delay cells that represent working memory. Many Delay cells fire during the cue and/or response epochs as well as the delay epoch; given their variable responses to the cue and response epochs, data analyses focused on the delay epoch. Unpaired t-test with Welch’s correction or one-way ANOVA or ordinary two-way ANOVA were employed to assess the effects of drug application on task-related activity or each single Delay cell. Two-tailed paired t-test t or repeated measures one-way ANOVA with Turkey’s multiple comparisons or repeated measures two-way ANOVA with Sidak’s multiple comparisons were employed to assess the effects of drug application on task-related activity for the population analysis. In the interest of brevity, figures often show the neurons’ preferred direction in comparison to just one non-preferred direction, the “anti-preferred” direction directly opposed to the neurons’ preferred direction. For Delay cells, the spatial tuning was assessed by comparing firing levels for the neuron’s preferred direction vs. its non-preferred directions. Quantification of spatial tuning was performed by calculating a measure of d’ using the formula:

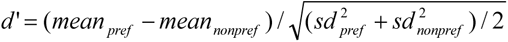

#### Cognitive behavior with systemic drug treatment

Rhesus monkeys (n=21, 10-33 years, 9 male, 12 female) were pretrained on the delayed response task, a test of visuospatial working memory performed with a manual response in a Wisconsin General Test Apparatus. A variable delay version was used, with 5 delays ranging from “0” seconds to the delay length that yielded chance performance for each animal, e.g. 0, 5, 10, 15 and 20sec. Delay lengths randomly varied over the 30 trials that made up a daily test session. Monkeys were tested twice a week, and tested for highly palatable food rewards, limiting the need for food regulation. This version of the task allows free movement that is optimal for studying aged monkeys’ cognitive performance. Delays were adjusted for each animal to achieve stable baseline performance of about 70% correct, leaving room for either improvement of impairment in performance. Monkeys were tested by an experimenter who was highly familiar with the normative behavior of each animal, but unaware of drug treatment conditions. Animals were rated for changes in sedation/agitation and aggression using 9 point rating scales.

Once stable baseline performance was achieved, the monkeys were administered either the KAT-II inhibitor, PF-04859989 (0.003-3.0 mg/kg, sc, 2 hrs before testing) ; the IDO inhibitor, INCB024360 (0.1 mg/kg, po 2 hrs before testing), or the KAT-II inhibitor, N-acetyl cysteine, (1.0-10.0 mg/kg, po, 2 hrs). All results were compared to vehicle control. Animals were required to return to stable baseline performance prior to receiving a subsequent dose of drug, with a minimum washout of at least 10 days. Statistical analyses employed Pearson’s r test of correlations between drug efficacy and age, and repeated-measures dependent T test with two-tailed significance.

### DATA AND CODE AVAILABILITY

The datasets supporting the current study are available from the corresponding author on request.

### KEY RESOURCES TABLE

#### Chemicals and antibodies

| REAGENT or RESOURCE | SOURCE | IDENTIFIER |
| --- | --- | --- |
| Goat anti-Rabbit IgG (H+L)<br>Highly Cross-Adsorbed<br>Secondary Antibody, Alexa<br>Fluor Plus 488 | ThermoFisher | Cat # A32731 |
| Goat anti-Chicken IgY (H+L)<br>Cross-Adsorbed Secondary<br>Antibody, Alexa Fluor Plus<br>647 | ThermoFisher | Cat # A32933 |
| Goat anti-Mouse IgG (H+L)<br>Cross-Adsorbed Secondary<br>Antibody, Alexa Fluor 594 | ThermoFisher | Cat# A-11005 |
| Anti-GFAP, NeuroMab clone<br>N206A/8 | NeuroMab/Antibodiesinc | Cat #75-240; RRID:<br>10672299 |
| Anti-MAP2 antibody | Abcam | Cat # ab5392; RRID:<br>AB_2138153 |
| AADAT Polyclonal antibody | Proteintech | Cat # 13031-1-AP; RRID<br>AB_2877904 |
| Hoechst 33342 | ThermoFisher | Cat #H23570 |
| Rabbit anti-kynurenic acid<br>polyclonal antibody | Abcam | Cat# ab37105,<br>RRID:AB_775944 |
| Kynurenic acid | Tocris Bioscience | Cat # 3694; CAS # 2439-02-3; PubChem CID # 52974250 |
| PF-04859989 | Sigma-Aldrich | Cat # 1122; CAS # 177943-33-8; PubChem CID #<br>329823792 |
| D-serine | Tocris Bioscience | Cat # 0226; CAS # 312-84-5;<br>PubChem CID # 71077 |
| PHA543613 | Tocris Bioscience | Cat # 3092; CAS # 1586767-92-1; PubChem CID # 56972222 |
| Antigen Unmasking Solution | Vector Labs | Cat # H-3300-250 |
| ProLong Gold Antifade Mountant | Invitrogen | Cat# P36930 |
| Biotin-SP (long spacer)<br>AffiniPure F(ab')<sub>2</sub> Fragment<br>Goat Anti-Rabbit IgG (H+L) | Jackson ImmunoResearch | Cat # 111-066-003<br>RRID: AB_2337966 |
| 3,3-diaminobenzidine tetrahydrochloride substrate system | Sigma Aldrich | D7304-1SET |
| Durcupan ACM, Aradite Embedding for Electron Microscopy | Electron Microscopy Sciences | Cat# 14040 |
| Vectastain Avidin-biotin peroxidase complex (ABC) kit | Vector Laboratories | AK-5000 |
| Osmium tetroxide (4% aqueous) | Electron Microscopy Sciences | Cat# 19140 |
| Normal Donkey Serum | Jackson ImmunoResearch | Cat# 017-000-121 |
| Bovine Serum Albumin | Sigma-Aldrich | Cat# A9467-100G |
| Triton X-100 | Sigma-Aldrich | Cat# X100-100ML |
| 10X Tris Buffered Saline (TBS) | Biorad | Cat# 1706435 |
| Hydrogen peroxide | J.T. Baker | Cat# 2186-01 |
| Uranyl Acetate | Electron Microscopy Sciences | Cat# 22400 |
| Propylene oxide | Electron Microscopy Sciences | Cat# 20401 |
| Ethanol | Decon Laboratories | Cat# 64-17-5 |
| Sodium phosphate monobasic | J.T. Baker | Cat# 3818-01 |
| Sodium phosphate dibasic anhydrous | J.T. Baker | Cat# 3828-05 |
| INCB024360 | Tocris Bioscience | Cat # 6007; CAS # 914471-09-3; PubChem CID # 11978742 |
| N-acetylcysteine | Tocris Bioscience | Cat # 7874; CAS #616-91-1; PubChem CID # 12035 |

#### Experimental Models: Organisms/Strains

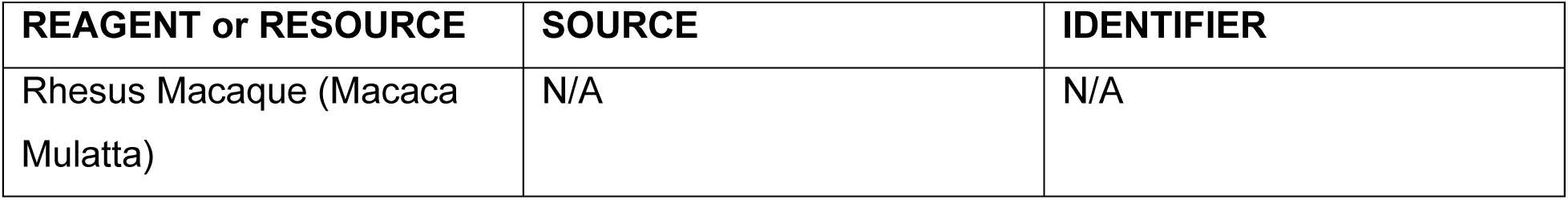

#### Software and Algorithms

| REAGENT or RESOURCE | SOURCE | IDENTIFIER |
| --- | --- | --- |
| MATLAB v. 2018b | MathWorks | https://www.mathworks.com/ |
| GraphPad Prism | GraphPad Software | https://www.graphpad.com/scientific-software/prism/ |
| SPSS Statistics | IBM | https://www.ibm.com/products/spss-statistics |
| CorelDRAW standard 2021 | CorelDRAW | https://www.coreldraw.com/ |
| Adobe Illustrator | Adobe | https://www.adobe.com/ |
| 3D Slicer | Slicer | www.slicer.org |

