## Supplemental FigureS1-5 for "Kynurenic acid inflammatory signaling expands in primates and impairs prefrontal cortical cognition"

**Figure S1- RORB is expressed in pyramidal cells in multiple layers in macaque dlPFC, including layer III.** RORB immunolabeling is prominently observed in rhesus macaque dlPFC layer III pyramidal cells, identified based on their canonical triangular-shaped soma morphology and well-defined apical dendrites (a-b). The methods for RORB immunohistochemistry have been described previously, and the RORB protein results for RORB are consistent with primate RNA-seq results. Scale bar: 20μm.

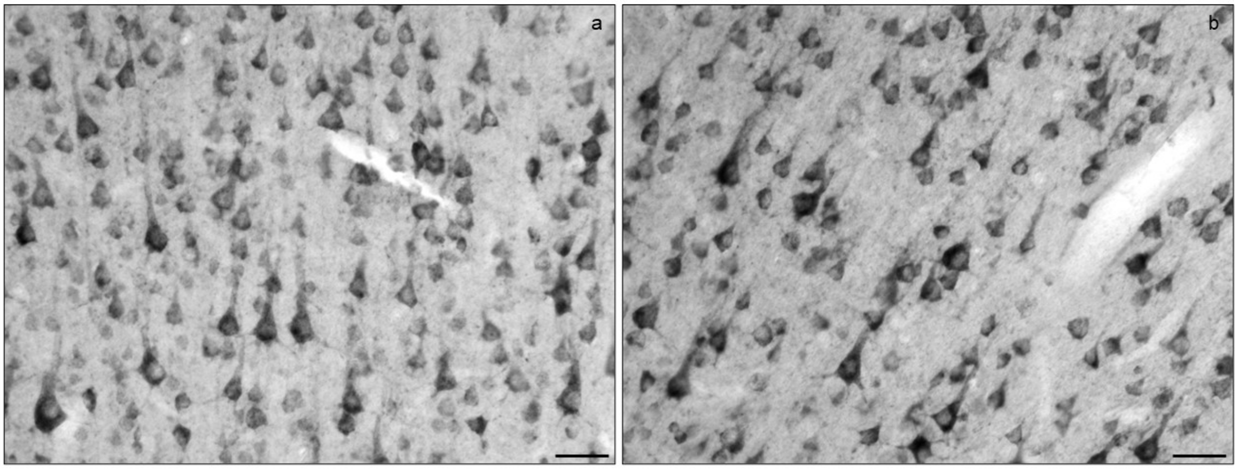

**Figure S2**- **Kynurenic acid immunolabeling by immunohistochemistry in aged macaque dlPFC.** High-magnification micrographs showing immunoperoxidase labeling for kynurenic acid in aged macaque dlPFC layer III expression in multiple cell-types, including neurons with pyramidal cell-like characteristics including triangular shaped cell bodies and labeling within well-defined apical dendrites (**a-d**). Kynurenic acid protein is also observed in the neuropil. Scale bars: 20μm. High magnification micrographs showing kynurenic acid immunoperoxidase immunolabeling in dlPFC white matter. The kynurenic acid immunolabeling is observed in the cell soma and extending along the radial processes of putative star-shaped astrocytes and bipolar oligodendrocytes. Scale bar: 20μm.

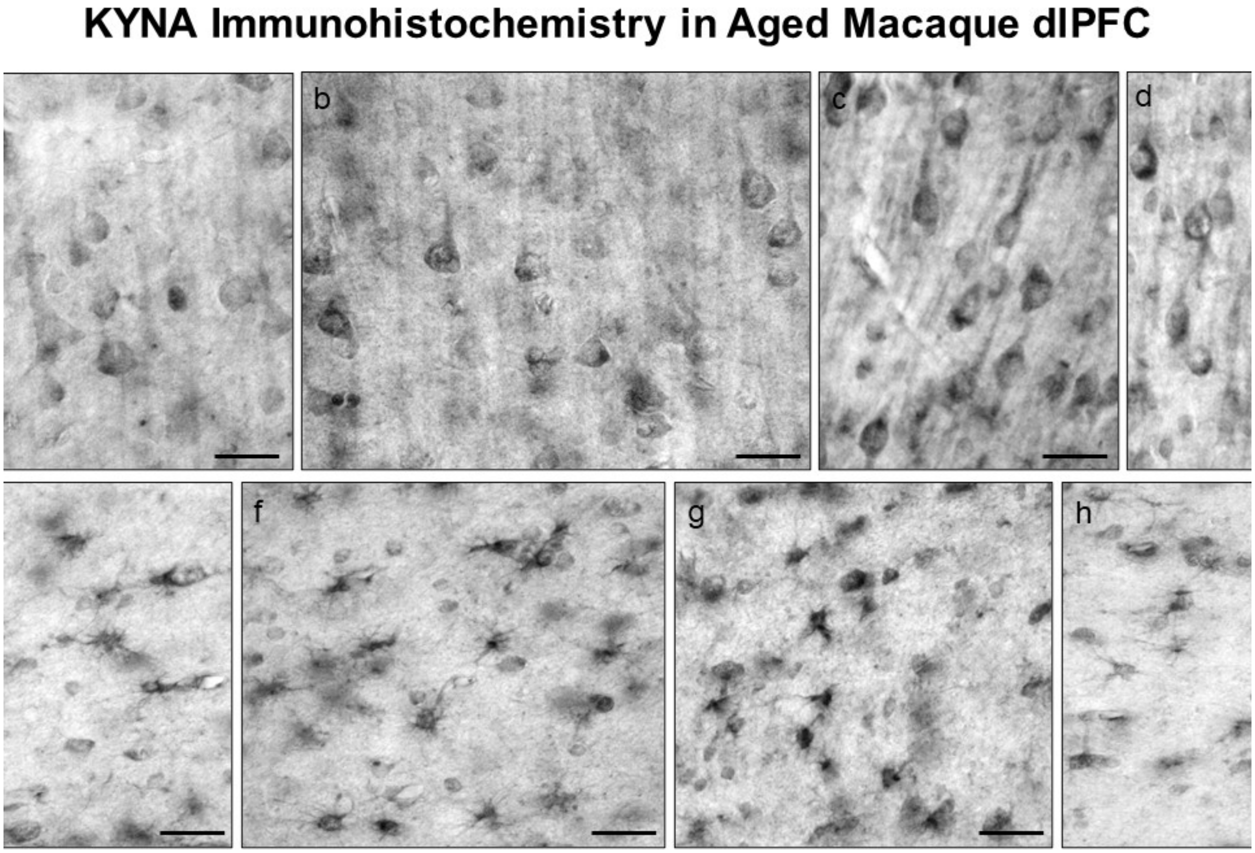

**Figure S3- Kynurenic acid is expressed in dendritic spines near the SER spine apparatus and astroglial leaflets in aged macaque dlPFC layer III.** Kynurenic acid immunolabeling (indicated by red arrowheads) is prominently expressed in dendritic spines (pseudocolored yellow), frequently near the calcium-storing smooth endoplasmic reticulum (SER) spine apparatus (pseudocolored pink; **a**). Additional examples of kynurenic acid immunolabeling on the plasma membrane of astrocytic leaflets (pseudocolored in green), within PAPs (**b-f**) near the excitatory glutamatergic synapse. All dendritic spines receive axospinous Type I asymmetric glutamatergic-like synapses, which are between arrows. As, astrocyte; Ax, axon; Mit, mitochondria; Sp, dendritic spine. Scale bars, 200 nm.

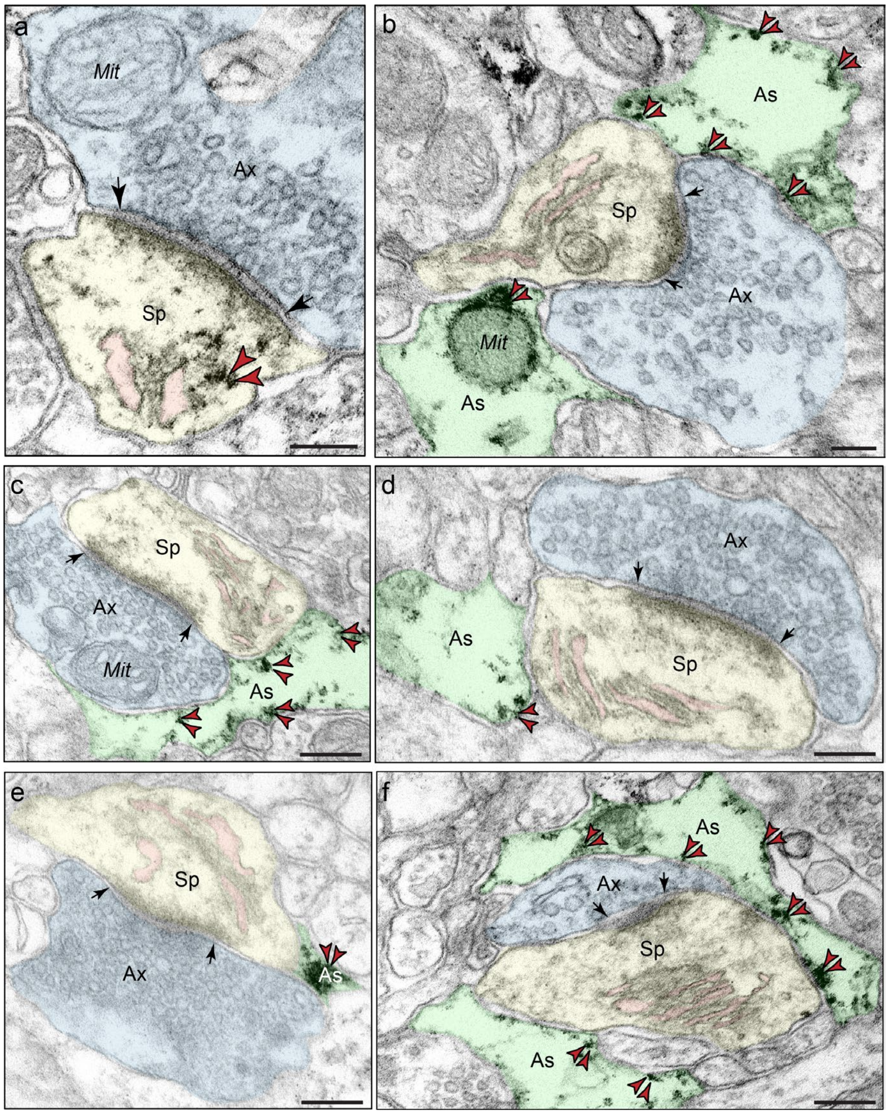

**Figure S4- KYNA reduced delay firing and spatial tuning of young macaque dlPFC Delay cells.**

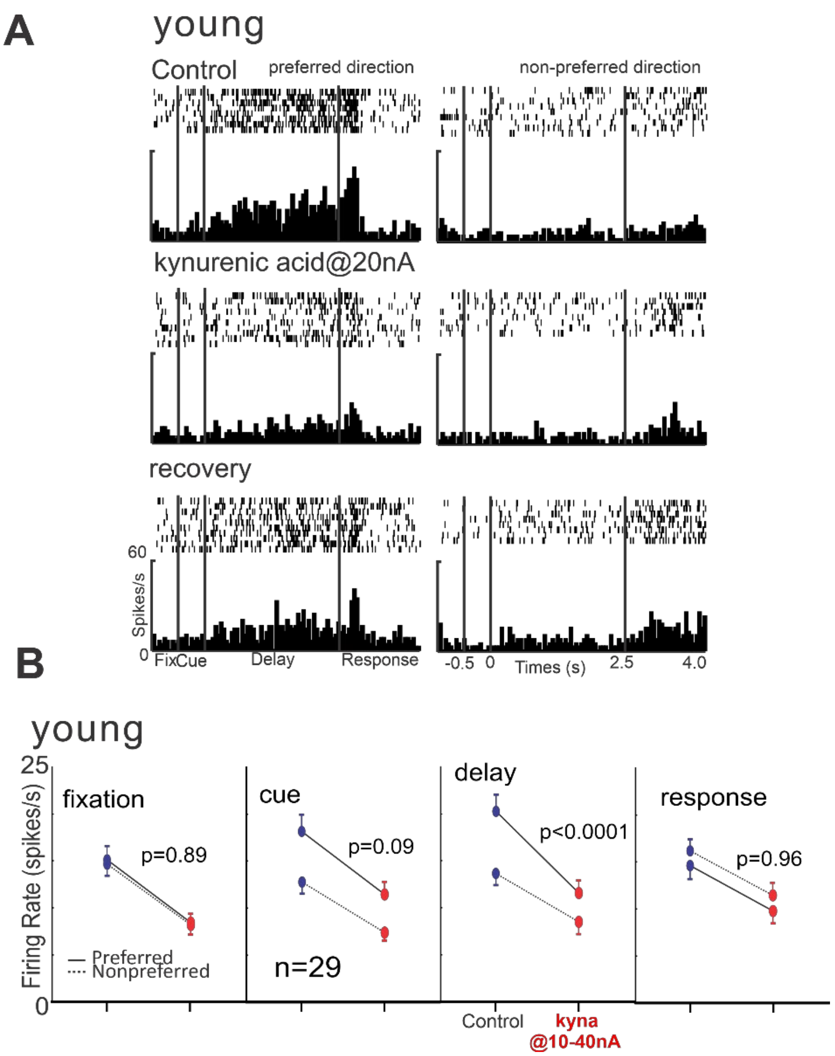


**A.** An example neuron is shown that iontophoresis of KYNA markedly reduced delay firing for the neuron's preferred direction, and the firing was partially restored when KYNA application was terminated (recovery). Raster and histogram data shown for the preferred and nonpreferred directions. **B**. KYNA reduced the delay firing and spatial tuning of Delay cells (n=29) from an young monkey. In contrast, there was no direction-specific effect of KYNA on either the fixation-, cue- or response-related firing.

**Figure S5- KYNA reduced delay firing and spatial tuning of middle-aged macaque dlPFC Delay cells.**

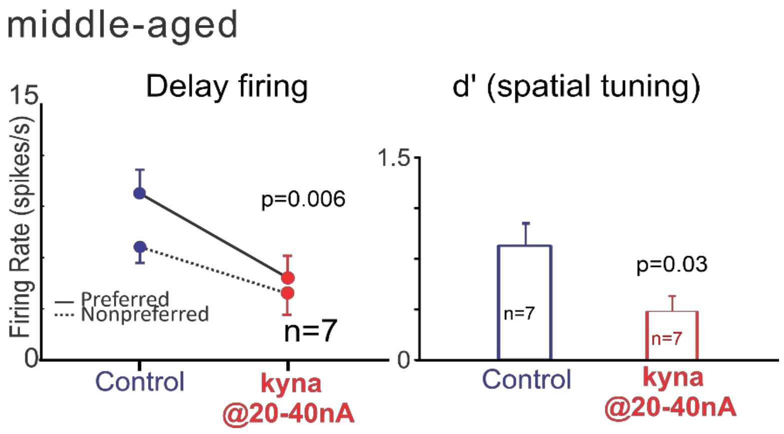


KYNA reduced the delay firing (2-way ANOVA-R, $F(1, 6) = 16.59$, $p=0.0066$) and spatial tuning (d': paired t test, $t=2.741$, $p=0.0337$) of Delay cells ($n=7$) from an middle-aged monkey.
